# Altered dynamical integration/segregation balance during anesthesia-induced loss of consciousness

**DOI:** 10.1101/2022.05.20.492794

**Authors:** Louis-David Lord, Timoteo Carletti, Henrique Fernandes, Federico E. Turkheimer, Paul Expert

**Affiliations:** Department of Psychiatry, University of Oxford, UK; Institut Méditerranéen de Recherches Avancées (IMéRA), Aix-Marseille University, France; Department of Mathematics and Namur Institute for Complex Systems (naXys), University of Namur, Belgium; Centre for Music in the Brain, Department of Clinical Medicine, Aarhus University, Denmark; Department of Neuroimaging, Institute of Psychiatry, Psychology and Neuroscience, King’s College London, UK; Global Business School for Health, University College London, UK

## Abstract

In recent years, brain imaging studies have begun to shed light on the neural correlates of physiologically-reversible altered states of consciousness such as deep sleep, anesthesia, psychedelic experiences. The emerging consensus is that normal waking consciousness requires the exploration of a dynamical repertoire enabling both global integration i.e. long-distance interactions between brain regions, and segregation, i.e. local processing in functionally specialized clusters. Altered states of consciousness have notably been characterized by a tipping of the integration/segregation balance away from this equilibrium. Historically, functional MRI (fMRI) has been the modality of choice for such investigations. However, fMRI does not enable characterization of the integration/segregation balance at sub-second temporal resolution. Here, we investigated global brain spatiotemporal patterns in electrocorticography (ECoG) data of a monkey (*Macaca fuscata*) under either ketamine or propofol general anaesthesia. We first studied the effects of these anesthetics from the perspective of band-specific synchronization across the entire ECoG array, treating individual channels as oscillators. We further aimed to determine whether synchrony within spatially localized clusters of oscillators was differently affected by the drugs in comparison to synchronization over spatially distributed subsets of ECoG channels, thereby quantifying changes in integration/segregation balance on physiologically-relevant time scales. The findings reflect global brain dynamics characterized by a loss of long-range integration in multiple frequency bands under both ketamine and propofol anaesthesia, most pronounced in the beta (13-30Hz) and low-gamma bands (30-80Hz), and with strongly preserved local synchrony in all bands.

## Introduction

Brain activity is characterized by the dynamic exploration of a diverse and flexible repertoire of functional brain configurations over time (1, 2). These explorations of the brain’s repertoire notably enable both global integration, i.e. long-distance interactions between brain regions, and local processing in functionally specialized clusters, i.e. segregation (3-5). The simultaneous occurrence of integration and segregation, a hallmark of complex systems displaying emerging properties (6, 7), is thought to lead to activity patterns of sufficiently high neural complexity to underlie normal, waking consciousness (8, 9).

It follows that events, be they pathological or pharmacological, which disrupt the integration/segregation balance should in turn modulate one’s behavior and/or subjective experience. Notably, functional MRI (fMRI) dynamic connectivity has been investigated in different sleep stages, pharmacologically induced anesthesia in humans and during psychedelic experiences. The emerging consensus is that unconscious states are associated with a loss of functional integration, as dynamic explorations become limited to specific patterns dominated by rigid functional configurations tied to the anatomical connectivity (9-13). On the other hand, psychedelic experiences, such as those induced by taking LSD or psilocybin, have been associated with increased global integration and dynamical explorations loosely constrained by the structural connectome (14-18). It is however important to note that, because fMRI allows for brain activity measurements on the order of seconds, the aforementioned studies have not yet determined how the integration/segregation balance is modulated on a sub-second scale in physiologically-reversible states of altered consciousness, such as general anesthesia.

Here, we investigated global brain spatiotemporal patterns in electrocorticography (ECoG) data of monkeys under either ketamine or propofol general anesthesia from the publicly available NeuroTycho project (19). While administration of both ketamine and propofol induces a rapid decrease in level of consciousness, the respective pharmacodynamics of these drugs markedly differs. Ketamine acts primarily as a NMDA receptor antagonist whereas propofol’s main target is the GABA_A_ receptor on which it has a positive allosteric effect. ECoG is an invasive imaging modality in which an electrode array is applied directly on the cortical surface. This allows for brain activity measurements on significantly faster timescales than fMRI. For example, synchronization in the beta (13-30 Hz) and low gamma (∼30-80Hz) frequency bands between neuronal populations is thought play a crucial role in the binding of information into conscious representations (20-23) and evidence shows it acts as a substrate for the emergence of functional networks (24). These links between high-frequency electrical oscillations and higher level cognitive entities cannot be directly investigated with fMRI due to insufficient temporal resolution.

In the present study, we first investigated the effects of ketamine and propofol from the perspective of band-specific synchronization behavior across the entire ECoG array, analyzing the collective behavior of ECoG channels treated as oscillators. We further sought to determine whether synchrony within spatially localized clusters of oscillators was differently affected by ketamine and propofol in comparison to synchronization over spatially distributed subsets of ECoG channels. This analysis hence enabled a quantification of the dynamics underlying the integration/segregation balance under anesthesia on physiologically-relevant timescales. The findings reflect global brain dynamics characterized by a loss of global integration in multiple frequency bands under both ketamine and propofol anesthesia, most pronounced in the beta (13-30Hz) and low-gamma bands (30-80Hz). Furthermore, whilst long-range synchronization was reduced during anesthesia relative to wakeful rest in those same frequency bands, synchrony within spatially localized electrode clusters remained strongly conserved for both anesthetic agents. Thus, despite their distinct pharmacodynamics, the effects of ketamine and propofol strongly converge on the novel measures of neural synchronization here applied to ECoG data.

## Methods

### Electrocorticography data acquisition and preprocessing

Neuroimaging data were obtained from the NeuroTycho dataset, an open-source set of 128-channel invasive electrocorticography (ECoG) recordings in a macaca fuscata monkey collected at the Riken Institute (Tokyo, Japan) (19). The ECoG array covered the left hemisphere only, with electrodes placed on both the lateral and medial brain surfaces. Data were recorded during both a resting-state baseline with eyes closed and during either propofol or ketamine-induced anesthesia in separate experiment. Each experiment was carried out least four days apart. During each experiment, the monkey was seated in a primate chair with both arms and head movement restricted. The ‘anesthesia’ condition was considered to have begun once the monkey failed to respond to sensory stimulation (i.e. stimulation of the nose with a cotton swab and unresponsiveness to have their forepaw touched). In the propofol condition, the monkey was given a single bolus of intravenous propofol (5.2 mg/kg), until loss of consciousness was observed. For the ketamine condition, a single bolus of intramuscular ketamine (5.1 mg/kg) was administered. For either drug, no maintenance anesthesia was given following initial induction.

ECoG data were acquired at a 1000 Hz sampling rate. As is common practice in electrophysiological studies, the quality of ECoG signals was assessed by visually inspecting the power spectra of individual ECoG channels, and no channels were discarded on this basis. The data were notch filtered at 50Hz to account for electrical line noise in Japan. While we did not downsample the data, we re-binned the timecourses into 100ms windows in order for our measures of inter-areal synchrony (see sections below) to operate on physiologically meaningful timescales. To ensure a consistent number of time points across all experimental conditions (ketamine/propofol and baseline), we considered the first 5000 time points of each experimental run, which yields a 5000*100ms = 500 sec (8.33 min) recording interval.

### Calculation of band-specific phase angles

We bandpassed each ECoG channel timecourse for the frequency-band of interest: *δ* = 0-4Hz; θ = 4-8Hz; α = 8-13Hz; β = 13-30Hz and γ = 30-80Hz. The bandpassed filtered data were normalized by removing the mean of each channel and dividing it by its standard deviation. For each ECoG channel *n*, we compute its analytic signal 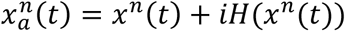 using the Hilbert transform *H*(*x*(*t*)). We then extract the instantaneous phase 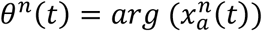. All analyses were performed using Matlab (MathWorks, Inc.).

We visualized the time-evolution of the phase angles for each ECoG channel in different frequency bands for both experimental conditions. For visualization purposes, only the first 500 time steps were plotted. To illustrate this output, the time-evolution of phase angles *θ*^*n*^(*t*) for ketamine, propofol and the respective wakeful rest baseline are shown in Figure 1 for the β-band (13-30Hz):

**Figure 1.**
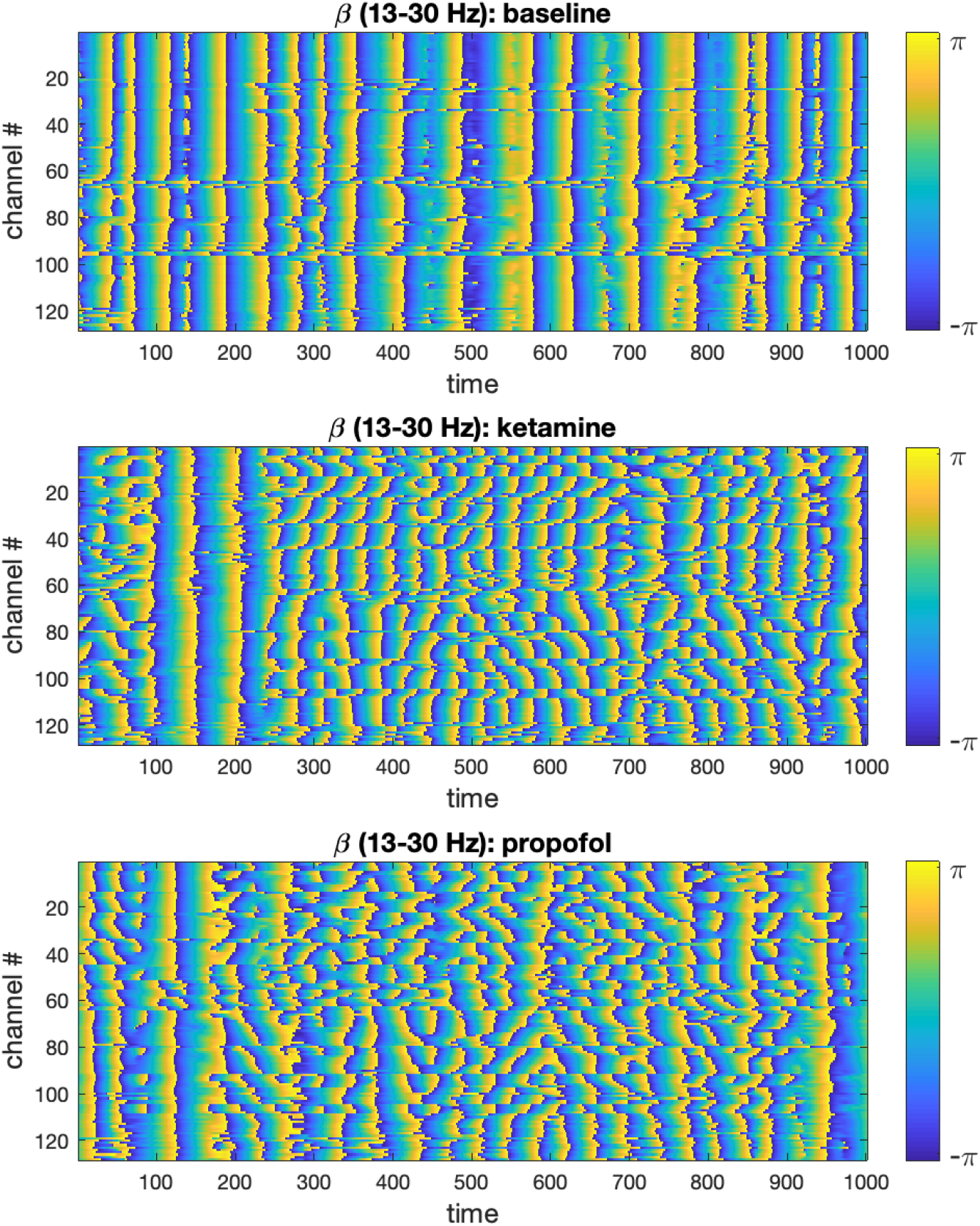
Time-evolution of phase angles *θ* derived from the ECoG signal via Hilbert transformation in the β-band is plotted for 1000 time steps (binned 100ms intervals) in the baseline, ketamine and propofol states. It can be qualitatively observed that for both drugs, the phase angles are on average less aligned at any given time point. We only show the β -band here for concision and to illustrate the methodology, but this same analysis step was performed for the other frequency bands under study (δ, α, γ) to enable statistical comparison (using all 5000 time steps).

### Analysis of global synchronization of ECoG channels

For each of the aforementioned frequency bands, we computed the Kuramoto order parameter (OP) (25) of the whole set of recording electrodes (n = 128 ECoG channels) at each time point (t = 5000):

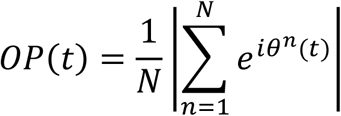

This measure enables us to quantify the extent of the global desynchronization induced by ketamine and propofol anesthesia relative to the wakeful rest baseline. The time-evolution of the OP in each frequency band for the anesthesia and baseline conditions were plotted, see Figure 2. Differences in the average value of the OP were compared between the baseline and anesthesia conditions by using paired t-tests (undirected) with Bonferroni correction for multiple comparisons.

**Figure 2.**
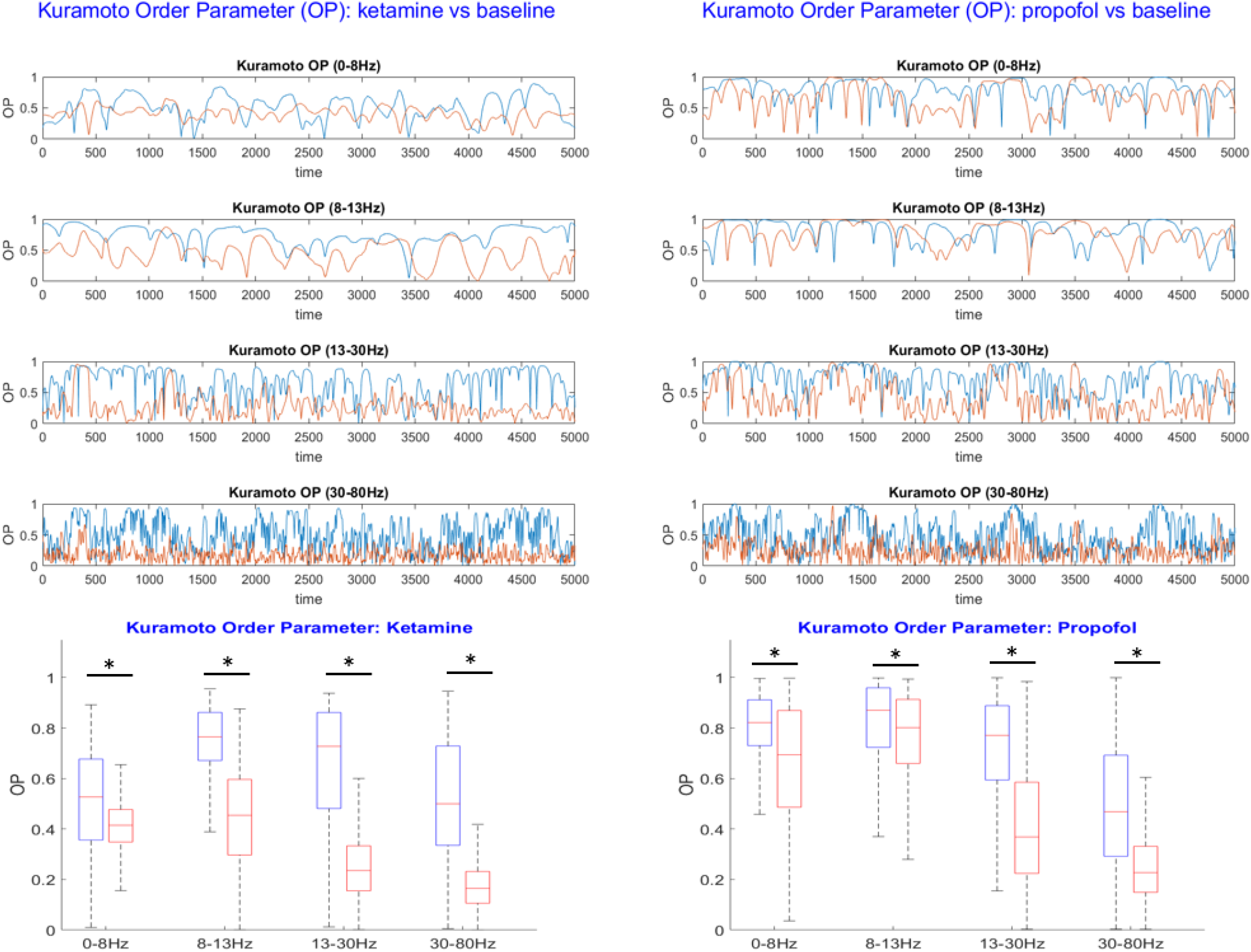
Top: The Kuramoto order parameter (OP) is plotted against time for both the anesthesia condition (red) and the wakeful rest baseline (blue) in each frequency band under study. Results from the ketamine anesthesia are shown on the lefthand side, and results for the propofol anesthesia on the right. Both ketamine and propofol induce a strong and statistically significant desynchronization of brain activity across the entire array of ECoG channels, persistent over all frequency bands under study. The largest effects of anesthetics on global synchrony were observed in the β (13-30Hz) and γ (30-80Hz) frequency bands, respectively. Bottom: Boxplots showing the mean OP over time ± standard deviation for the baseline (blue) and anesthesia conditions (red). Statistically significant reductions in global synchrony were observed across all frequency bands for both ketamine (left) and propofol (right). The largest effects on global synchrony were observed in the β (13-30Hz) and γ (30-80Hz) frequency bands.

### Comparison of local vs distributed cluster synchrony

We then shifted our attention to studying the brain dynamics from the perspective of clusters of electrodes seen as oscillators. To do so, we used the Matlab implementation of k-means clustering, the kmeans function, to identify groups of ECoG channels based on their geometry, clustering together *spatially local* channels, for: *k*= 13, 14, 15. These *k*-values ensure that the number of electrodes per cluster is small enough to obtain well localized clusters that contain a non-trivial number of electrodes. To deal with the latter point, we introduced a constraint in the *k*-mean algorithm so that the smallest possible number of electrodes in a given cluster was n =4. To account for the randomness involved in k-means initialization and ensure replicability of our findings, we performed 100 iterations of the k-means algorithm in our analyses. A typical exemplar of community assignments obtained for *k* = 14 is shown in Figure *2*.a.

For each cluster of electrodes C, we calculated the internal synchrony *φ*_*C*_ at each time step *t*:

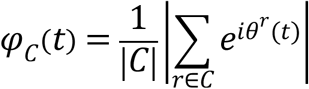

where |C| denotes the size of the cluster under scrutiny. Local synchrony was calculated for each frequency band on the clusters of ECoG channels defined by the k-means clustering algorithm, i.e. spatially compact clusters, for all k values (*k* = 13, 14, 15). This process was repeated for each of the 100 k-means iterations.

Distributed synchrony was calculated for each frequency band on the same number of clusters k as local synchrony, but with *random* electrode assignments to the clusters, see Figure *2*.b for an illustration. To ensure the robustness of this approach over a large number of randomly assigned electrode clusters, we computed local synchrony on 100 distinct sets of distributed clusters for *k* = 13, 14 and 15 respectively. For consistency, we introduced the same constraint as with the k-means, ensuring that the minimum size of any distributed cluster was *n* = 4 ECoG channels. As expected, randomly assigning the electrodes to k clusters statistically significantly increased the average Euclidian distance between electrodes in a given cluster compared to the corresponding k-means algorithm output, thus yielding *spatially distributed* cluster. See Figure *3* for a typical local and distributed electrodes and Figure 4 for the statistics on the spatial localization of the local and distributed clusters.

**Figure 3.**
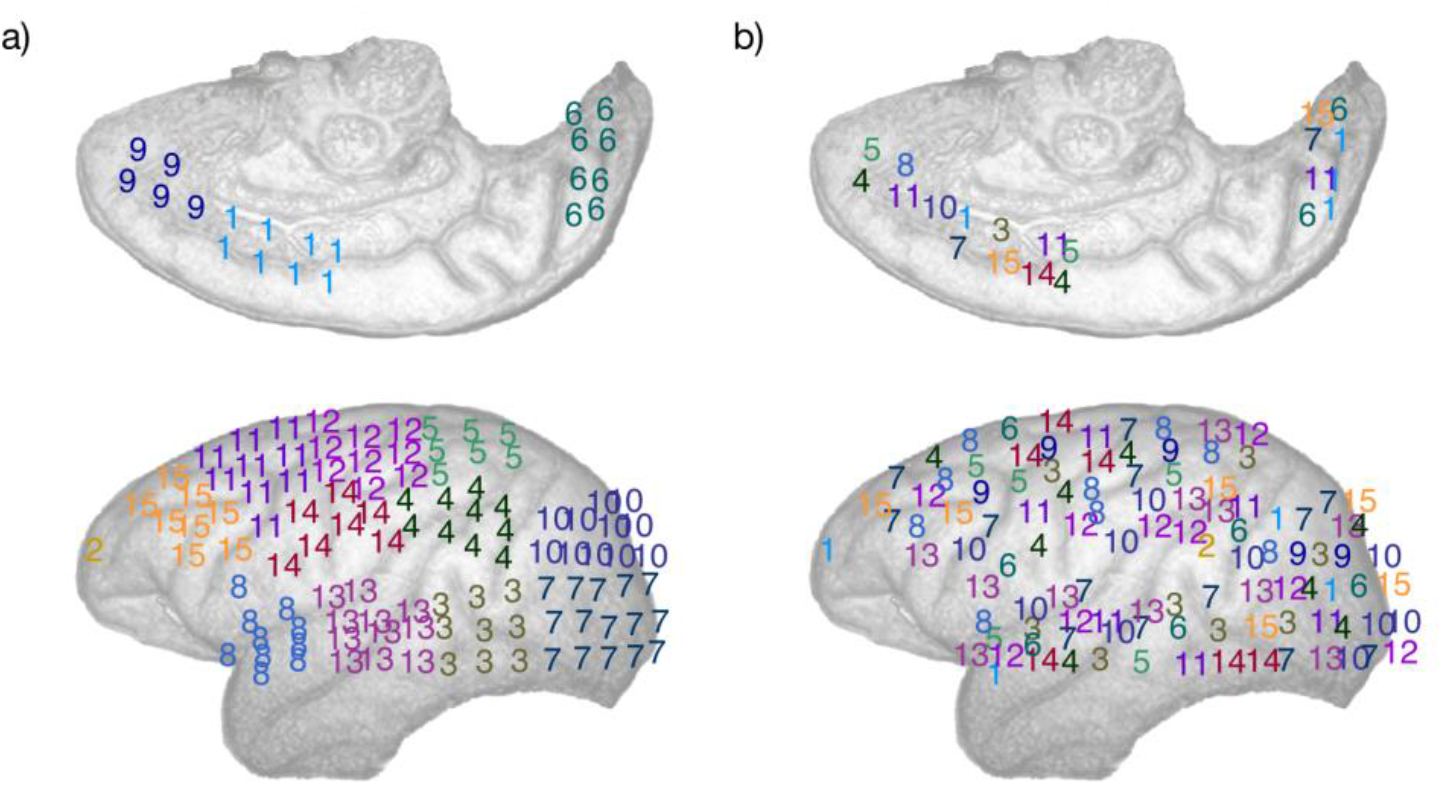
Schematic representation of the electrodes array implemented in the left cortical surface of the monkey, inside surface on the top row, outside surface on the bottom row. The electrodes are numbered according to: a) A typical spatially local clustering for the electrodes using k-means clustering with k=14. b) A typical spatially distributed clustering from random electrodes assignment for k=14. See Methods: Comparison of local vs distributed cluster synchrony for details.

**Figure 4.**
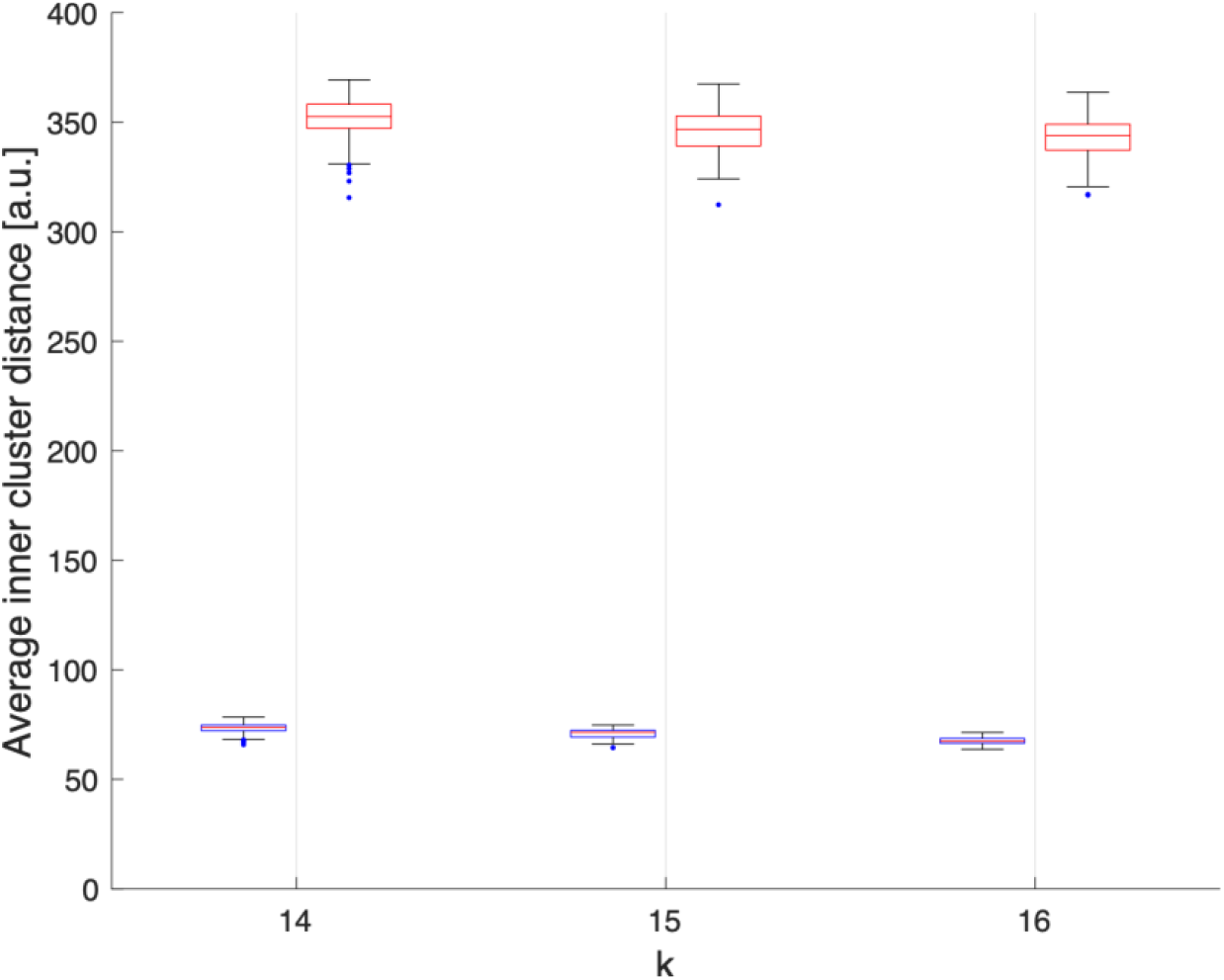
Average Euclidean distance, in arbitraty units, between electrodes within each clusters obtained assuming *k=14, 15 or 16* over 100 realisations. We can observe that electrodes grouped by k-means clustering are spatially compact (blue), while the electrodes grouped randomly are spatially delocalized (red).

For each frequency band and cluster number *k*, we compared the effects of ketamine and propofol anesthesia on the internal synchrony of local vs distributed electrode clusters using repeated paired t-tests with Bonferroni correction for multiple comparisons. Thus, the statistical significance of pre vs post-drug effect on cluster synchrony were separately calculated for 100 local cluster assignments, and 100 distributed cluster assignments respectively using a Bonferroni-adjusted significance threshold of α = 5*10^−4^.

## Results

### Ketamine and propofol induce global, frequency band-specific desynchronization of brain oscillations

In each frequency band under study, we find that both ketamine and propofol induce a global desynchronization of brain activity across the entire array of ECoG channels, as indicated by significant decreases in the Kuramoto order parameter (OP) of the system in all frequency bands considered, see Figure 4. For ketamine, mean percent decreases in OP during anesthesia relative to the wakeful rest baseline were: -21.3% (0-8Hz); -40.9% (8-13Hz); -59.1% (13-30Hz); -65.6% (30-80Hz) (all p-vals < 10^−200^). For propofol, mean percent decreases in OP during anesthesia relative to the wakeful rest baseline were: - 16.7% (0-8Hz); -5.9% (8-13Hz); -41.8% (13-30Hz); -49.1% (30-80Hz) (all p-vals < 10^−40^). We note that for both drugs, the largest effects on global synchrony were observed in the β (13-30Hz) and γ (30-80Hz) frequency bands. Whilst still statistically significant, the weakest effect was observed for propofol in the α band (8-13Hz).

### Differential effects of general anesthetics on synchronization spatially localized vs distributed electrode clusters

Because the Kuramoto order parameter (OP) considers the synchronization behavior of the entire electrode array, it does not allow to infer any distinction between local and distributed effects of the drugs on brain dynamics. We therefore followed up the initial analysis by comparing synchronization behavior of spatially localized and spatially distributed clusters of electrodes; the latter requiring long-range synchronization while the former does not. The internal synchrony, i.e. the local order parameter, of a given cluster at a given time step is denoted as *φ*_*C*_(*t*) We provide a visualization of this analysis in Figure *5* below; changes the time-evolution of *φ*_*C*_ are plotted for 14 clusters derived from beta-band activity (13-30 Hz) under the different experimental conditions. A marked reduction in *φ*_*C*_ is observed in the distributed clusters of electrodes for both ketamine and propofol anesthesia, but *φ*_*C*_ remains markedly unchanged between the baseline and anesthesia conditions for the spatially localized clusters.

**Figure 5.**
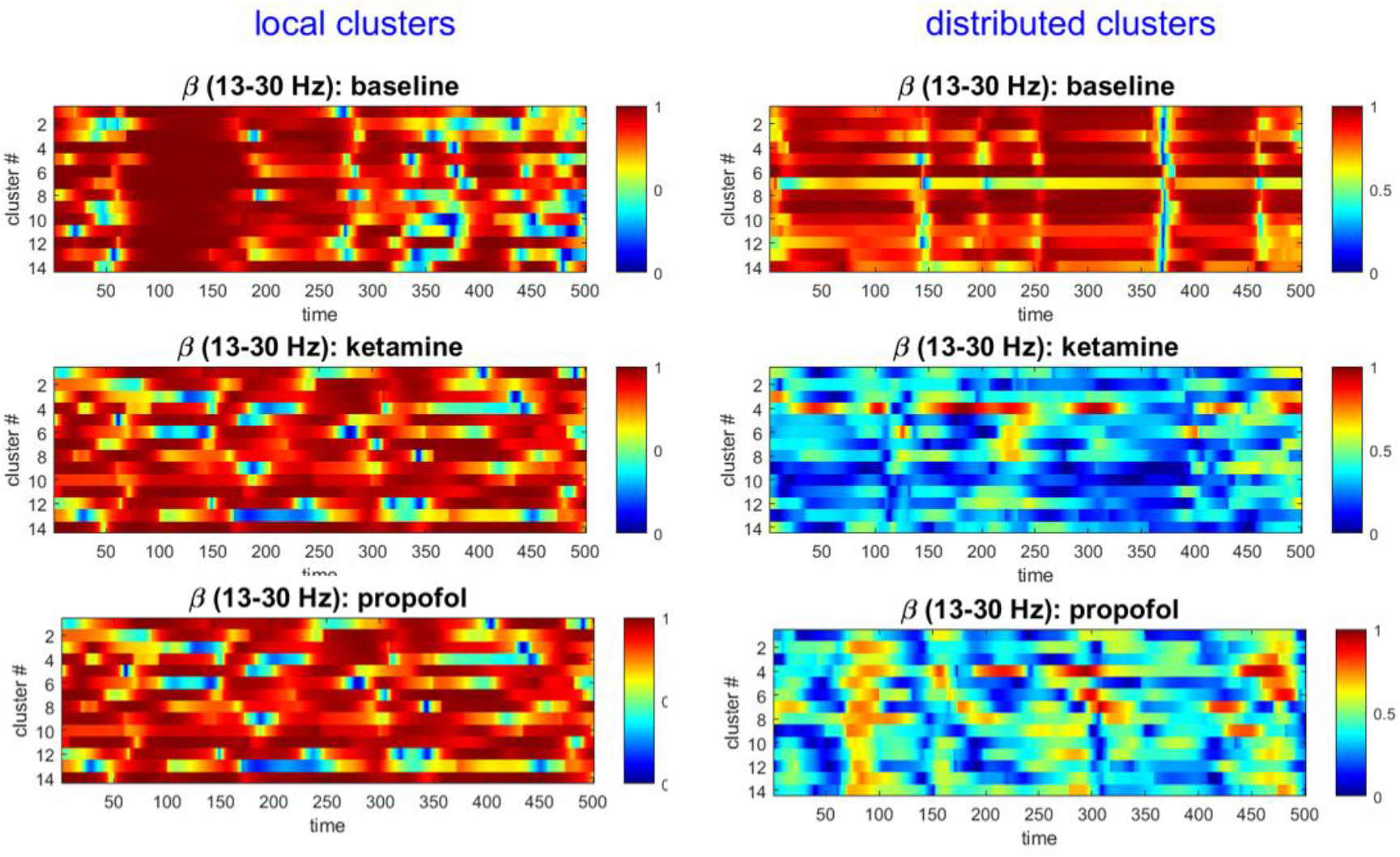
The local cluster synchrony *φ*_*C*_ is plotted against time for the first 500 time-steps (binned 100ms intervals) in the baseline, ketamine and propofol states for *k*=14 clusters that are either spatially localized (lefthand side) or spatially distributed (righthand side). We only show results from the clusters derived from the β -band here for concision and to illustrate the methodology, but this same analysis was performed for each frequency band under the study to enable statistical comparison (using all 5000 timesteps). Local cluster synchrony in β-band is preserved under both ketamine and propofol anesthesia. By contrast, distributed cluster synchrony is strongly decreased by each drug.

For both ketamine and propofol, synchrony was strongly maintained in spatially localized electrode clusters of ECoG channels, defined by k-means clustering; see Methods section, for *k* =14. For ketamine, synchronization in spatially localized clusters was not significantly different in any of the frequency bands at the Bonferroni-adjusted significance threshold (α = 5*10^−4^). The average p-values were: 0.618, 0.010, 0.019 and 0.383 across all 100 k-means iterations for the δ, α, β, and γ bands, respectively, see Figure *6* left column. Similarly, for propofol anesthesia, synchronization within local clusters was also preserved relative to baseline in all frequency bands and across all cluster assignments. The averaged p-values, obtained across all 100 k-means iterations, were: 0.77, 0.28, 0.74 and 0.76 for δ, α, β, and γ frequency bands, see Figure *7* left column.

**Figure 6.**
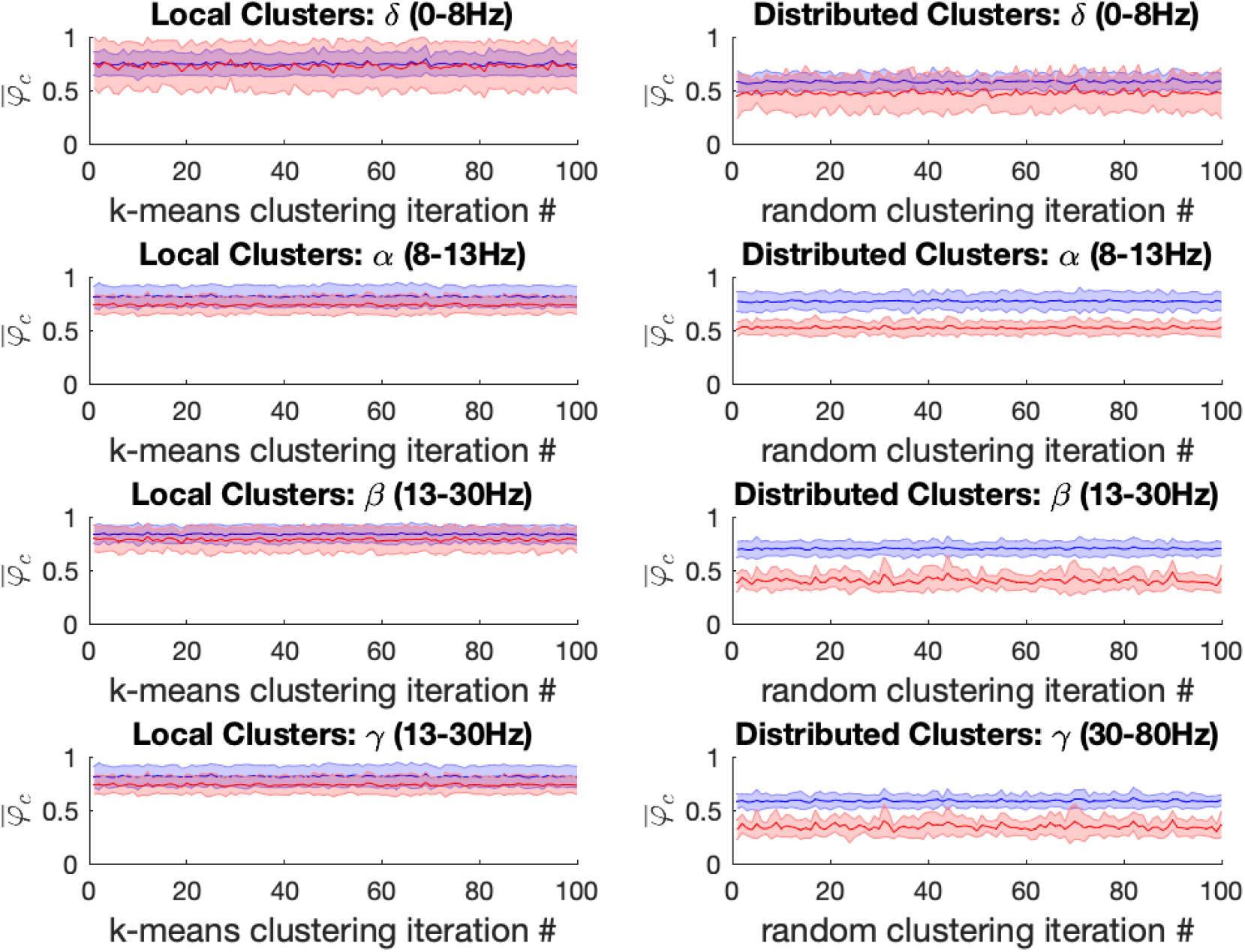
Left: For the ketamine (red) vs baseline (blue) conditions, the mean local cluster synchrony 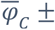 standard deviation is plotted for each of 100 k-means iterations (x-axis) in each of the four frequency bands under study for *k* = 14. Repeated within-condition t-tests failed to reach statistical significance at the Bonferroni-corrected threshold (α = 5*10^−4^) in any of the four frequency bands of interest. Right: Mean distributed cluster synchrony 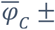 standard deviation is plotted for each of 100 iterations of the pseudo-random algorithm used for distributed cluster assignments (x-axis). Statistically significant reductions in distributed cluster synchrony at the Bonferroni-corrected significance threshold (α = 5*10^−4^) were observed following ketamine anesthesia for all (100 / 100) distributed cluster assignments in the α, β and γ bands, respectively. By contrast no significant differences in drug effects on distributed cluster synchrony were observed in the δ-band.

**Figure 7.**
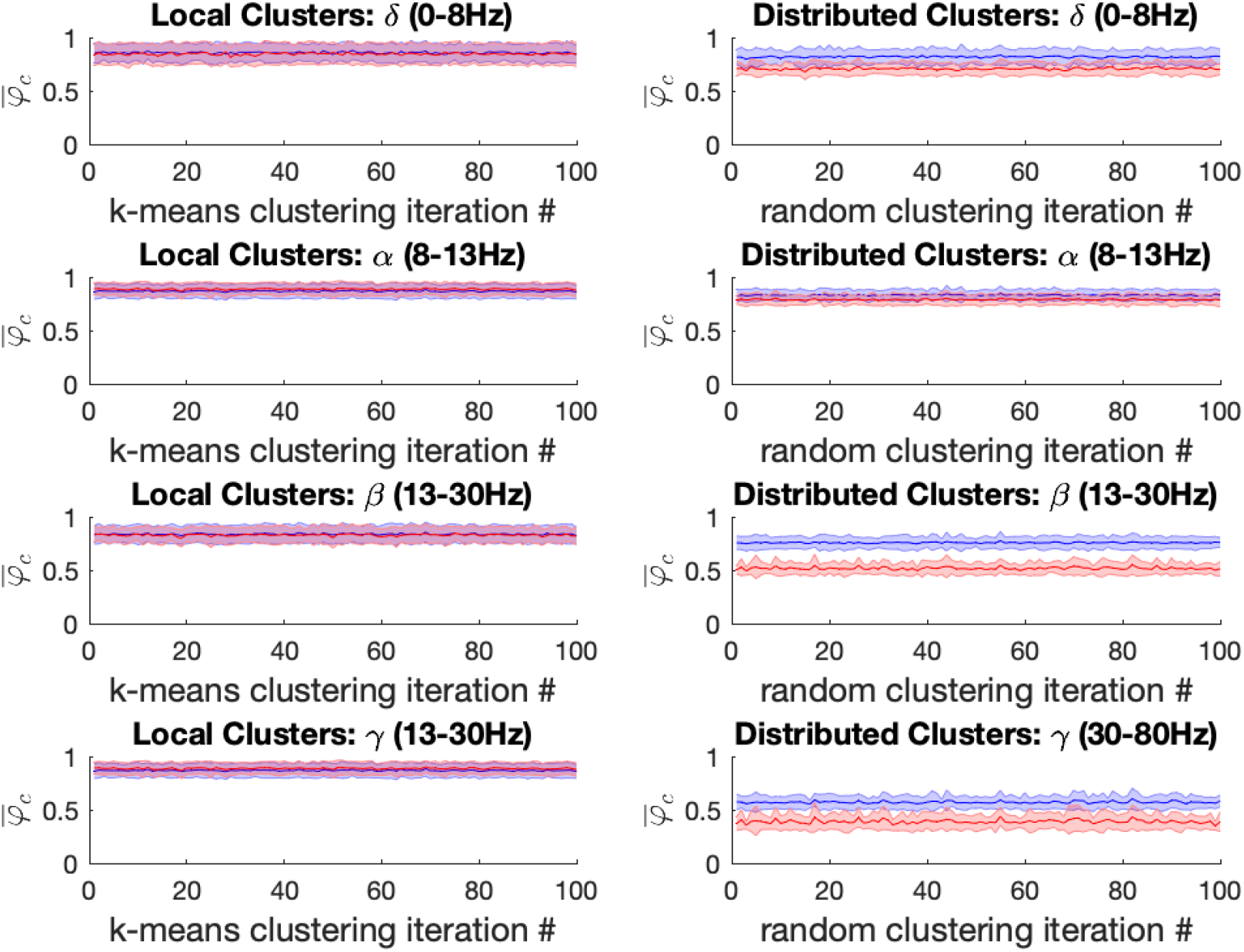
Left: For the propofol (red) vs baseline (blue) conditions, the mean local cluster synchrony 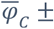 standard deviation is plotted for each of 100 k-means iterations (x-axis) in each of the four frequency bands under study for *k* = 14. Repeated within-condition t-tests failed to reach statistical significance at the Bonferroni-corrected threshold (α = 5*10^−4^) in any of the four frequency bands of interest. Right: Mean distributed cluster synchrony 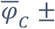 standard deviation is plotted for each of 100 iterations of the pseudo-random algorithm used for distributed cluster assignments (x-axis). Statistically significant reductions in distributed cluster synchrony at the Bonferroni-corrected significance threshold (α = 5*10^−4^) were observed following propofol anesthesia for all (100/100) distributed cluster assignments in the β and γ bands, respectively. By contrast only 6/100 significant differences in propofol’s effect on distributed cluster synchrony were observed in the α-band, while more mixed results were observed in the δ-band with 60/100 of distributed cluster assignments reaching significance.

Under ketamine anesthesia, synchronization in spatially distributed clusters was strongly and significantly reduced in every distributed cluster assignment (100/100) for the α, β, and γ bands, but no such reductions were observed in the δ band (0 / 100), see Figure *6* right column. The average p-vals were: 0.09, 3*10^−7^, 8*10^−7^ and 3*10^−6^ for δ, α, β, and γ frequency bands respectively (Bonferroni-adjusted significance threshold α = 5*10^−4^). Under propofol anesthesia distributed cluster synchrony was significantly reduced for every cluster assignment (100/100) in the β, and γ bands with average p-values of 1.1*10^−7^ and 2.7*10^−5^ respectively, see Figure *7* right column. Statistically significant reductions in distributed cluster synchrony were observed for 60/100 cluster assignments in the δ band (average p-val = 0.0012) but for only 6/100 cluster assignments in the α band (average p-val = 0.017).

Thus, for both ketamine and propofol, statistically significant reductions in distributed cluster synchrony, that are robust over cluster assignments, were observed in the β, and γ frequency bands. By contrast, synchronization in local clusters was not significantly impacted by either drug.

## Discussion

The integration/segregation balance is intrinsically linked to some of the most complex and fascinating phenomena of modern neuroscience; from the spontaneous emergence of resting-state networks to higher cognition and consciousness. Yet, relatively little is known about how changes in subjective experience relate to time-resolved changes in the integration/segregation balance. Here we applied novel analytical approaches to characterize changes in frequency band-specific synchronization patterns between clusters of ECoG channels in a monkey under general anesthesia, compared to a wakeful rest baseline. The effects of both ketamine and propofol anesthesia were studied across separate scanning session. We placed specific emphasis on investigating the dynamical integration/segregation balance during anesthesia-induced loss of consciousness by considering individual ECoG channels as oscillators.

Our findings describe a brain functional architecture characterized by a loss of global functional integration during both ketamine and propofol-induced loss of consciousness caused by a selective disruption in long-range synchrony, particularly in the β and γ frequency bands. On the other hand, synchronization in spatially localized clusters remained unaffected by both anesthetic drugs relative to a wakeful rest baseline in all frequencies under study. Furthermore, the results were highly consistent upon building electrode clusters of different sizes (k = 13, 14, 15) (see supplementary materials). These results were broadly consistent for both ketamine and propofol, hinting at a potential general mechanism underlying drug induced loss of consciousness. In other words, two drugs with markedly different pharmacodynamics led to the same outcome, both neurophysiologically on synchronization measures employed in this study, as well as behaviorally with regards to their general anesthetic properties.

The formation of dynamic links between brain regions via synchronization of oscillations in different frequency bands is one of the most plausible mechanisms for large-scale integration in the brain, and for the resulting binding of information into unified cognitive moments (21, 23, 24, 26). For both ketamine and propofol anesthesia, we found that long-range synchronization was most strongly disrupted in the beta (13-30Hz) and gamma (30-80Hz) frequency bands. Interestingly, oscillations in the beta frequency range have been shown to facilitate long-range interactions at the cortical level (21, 27, 28). Similarly, several lines of evidence have implicated gamma-oscillations in both long-range synchronization and as a neural correlate of conscious perception (20, 23, 29). Thus, our findings of strongly reduced long-range synchrony in these particular frequencies during anesthesia-induced loss of consciousness provides further evidence for the contribution of the beta and gamma bands toward the integration of information into conscious representations.

The present results are also in agreement with prior studies which have used dynamical network analyses and related approaches to study physiologically-reversible changes in consciousness. The emerging consensus from this growing body of fMRI literature is that conscious wakefulness is characterized by global integration, (i.e. long-distance interactions between brain regions displaying scaling behaviour in time and space across a range of imaging modalities) (30, 31), as well as by the dynamic exploration of a diverse and flexible repertoire of functional brain configurations (32-34). Conversely, during anesthesia or deep sleep, long-range functional integration is reduced and the dynamic explorations become limited to a smaller subset of specific activity patterns, more tightly constrained by the structural connectome (11). Our results therefore extend the idea of a dynamical imbalance between the integration and segregation of information during altered consciousness to electrophysiological data at much faster timescales than fMRI investigations allow for.

The measures introduced in this study capture global changes in the brain’s spatiotemporal organization induced by ketamine and propofol without considering the precise localization these changes within the brain. Specifically, they demonstrate the potential sensitivity of time-resolved measures of system-wide integration/segregation balance to changes in consciousness level and/or contents. We suggest that related measures could be extended beyond anesthesia and remain sensitive to a range of conditions in which behavior and/or subjective experience is measurably altered, i.e. psychopharmacological effects, neuropsychiatric illness, sleep, disorders of consciousness, etc. For example, we would notably expect psychedelic drugs, e.g. psilocybin and LSD, to have opposite effects as general anesthetic drugs on the measures employed in this study, since some fMRI studies have shown that these compounds may actually increase long-range functional integration (15, 16, 35). An extension of this idea is that biomarkers for neurological and psychiatric disorders need not be necessarily based on changes in specific brain areas or circuits, but could also be reflected as altered global spatiotemporal patterns. For example, a dynamical functional connectivity analysis of fMRI data from patients with major depressive disorder has revealed abnormally stable patterns of long-range synchronization, consistent with various clinical features of depressive patients, including ruminative, slow, and monotonous thinking (36, 37). The present work lays a foundation for further studies in this direction.

In conclusion, we investigated changes in the dynamical balance between integration and segregation of information during anesthesia-induced loss of consciousness in ECoG data from a macaque monkey. The analysis was sensitive to the breakdown of global functional integration and long-range synchronization previously documented in fMRI investigations of physiologically reversible unconscious states. We also report that, in contrast to long-range synchronization, functional synchrony in spatially localized clusters remained highly conserved under general anesthesia. We therefore propose that time-resolved measures of synchronization behavior in clusters of oscillators may be sensitive to departures from normal waking consciousness in various conditions and be clinically useful.

## Acknowledgements

L.D. Lord, T. Carletti, H. Fernandes and P. Expert conducted part of the study as Fellows of the Institut Méditeranéen d’Etudes et de Recherches Avancées (IMeRA).

## Supplementary Material

In this Appendix, we report the results of our analysis obtained on clusters of size k=13 and 15 for both Ketamine and Propofol. The results are consistent and robust with the k=14 case in the main text.

Figures *Supplementary 1* and Supplementary 2 show the mean synchrony, with standard deviation, for the baseline and Ketamine conditions in the local cluster (left) and the distributed clusters (right). Figures *Supplementary 3* and *Supplementary 4* show the mean synchrony, with standard deviation, for the baseline and Propofol conditions in the local cluster (left) and the distributed clusters (right). Tables *Supplementary 5* and *Supplementary 6* recapitulate the average p-values for the significance of the difference in the mean between the baseline and Ketamine conditions across all 100 electrode assignments for the local and distributed clusters respectively. Significance is attained at the Bonferroni corrected threshold (α = 5*10^−4^) only for the distributed cluster in the α, β and γ, consistently across of cluster numbers. Similarly, tables Supplementary 7 and *Supplementary 8* show the same information for the difference in synchrony between the baseline and propofol conditions. Significance is attained at the Bonferroni corrected threshold (α = 5*10^−4^) only for the distributed cluster in the β and γ, consistently across of cluster numbers.

**Supplementary 1.**
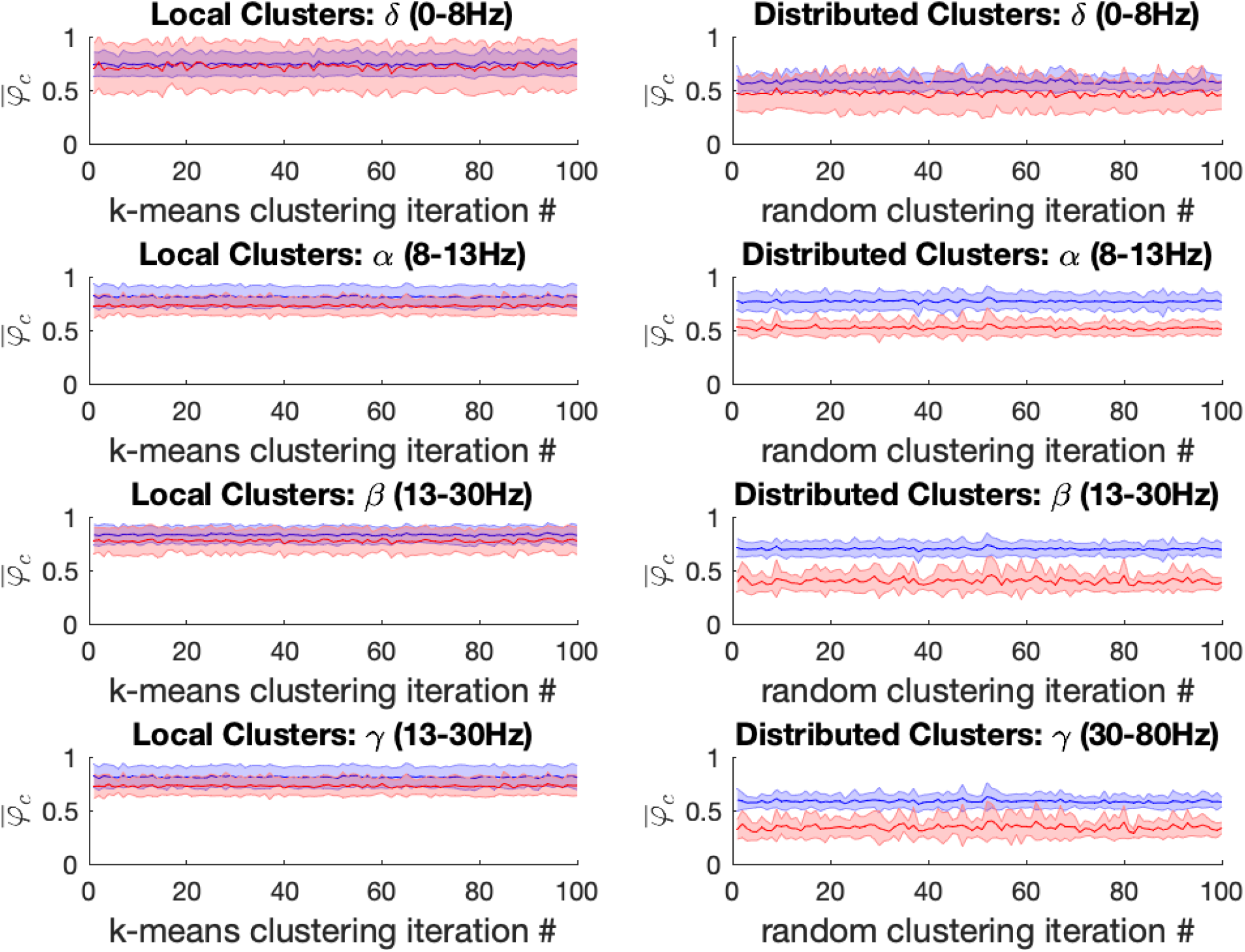
Left: For the ketamine (red) vs baseline (blue) conditions, the mean local cluster synchrony 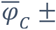 standard deviation is plotted for each of 100 k-means iterations (x-axis) in each of the four frequency bands under study for k = 13. Repeated within-condition t-tests failed to reach statistical significance at the Bonferroni-corrected threshold in any of the four frequency bands of interest. Right: Mean distributed cluster synchrony 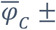 standard deviation is plotted for each of 100 iterations of the pseudo-random algorithm used for distributed cluster assignments (x-axis). Statistically significant reductions in distributed cluster synchrony at the Bonferroni-corrected significance threshold (α = 5*10^−4^) were observed following ketamine anesthesia for all (100 / 100) but one distributed cluster assignments in the α, β and γ bands, respectively. By contrast no significant differences in drug effects on distributed cluster synchrony were observed in the δ-band.

**Supplementary 2.**
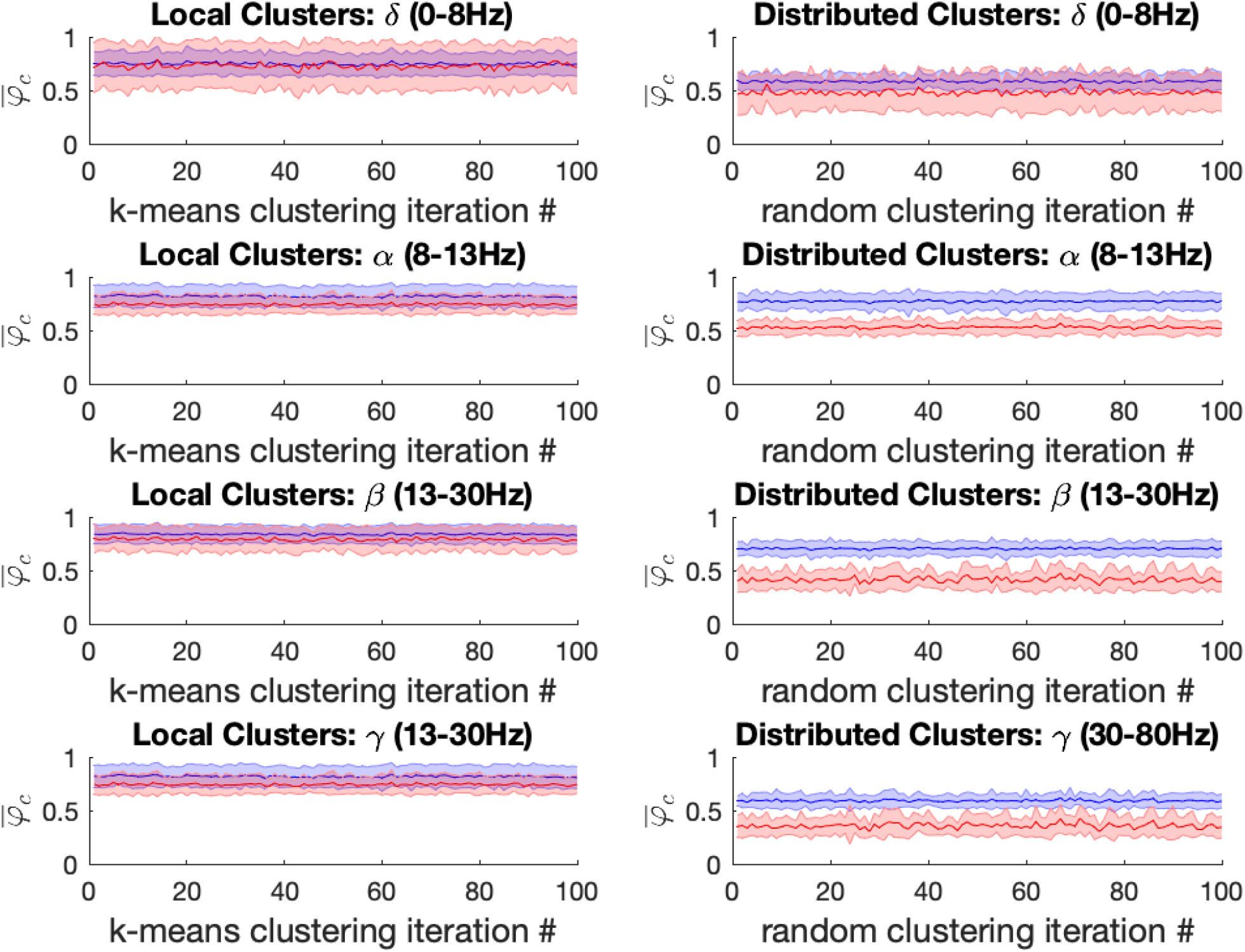
Left: For the ketamine (red) vs baseline (blue) conditions, the mean local cluster synchrony 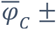 standard deviation is plotted for each of 100 k-means iterations (x-axis) in each of the four frequency bands under study for k = 15. Repeated within-condition t-tests failed to reach statistical significance at the Bonferroni-corrected threshold (α = 5*10^−4^) in any of the four frequency bands of interest. Right: Mean distributed cluster synchrony 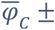 standard deviation is plotted for each of 100 iterations of the pseudo-random algorithm used for distributed cluster assignments (x-axis). Statistically significant reductions in distributed cluster synchrony at the Bonferroni-corrected significance threshold (α = 5*10^−4^) were observed following ketamine anesthesia for all (100 / 100) distributed cluster assignments in the α, β and γ bands, respectively. By contrast no significant differences in drug effects on distributed cluster synchrony were observed in the δ-band.

**Supplementary 3.**
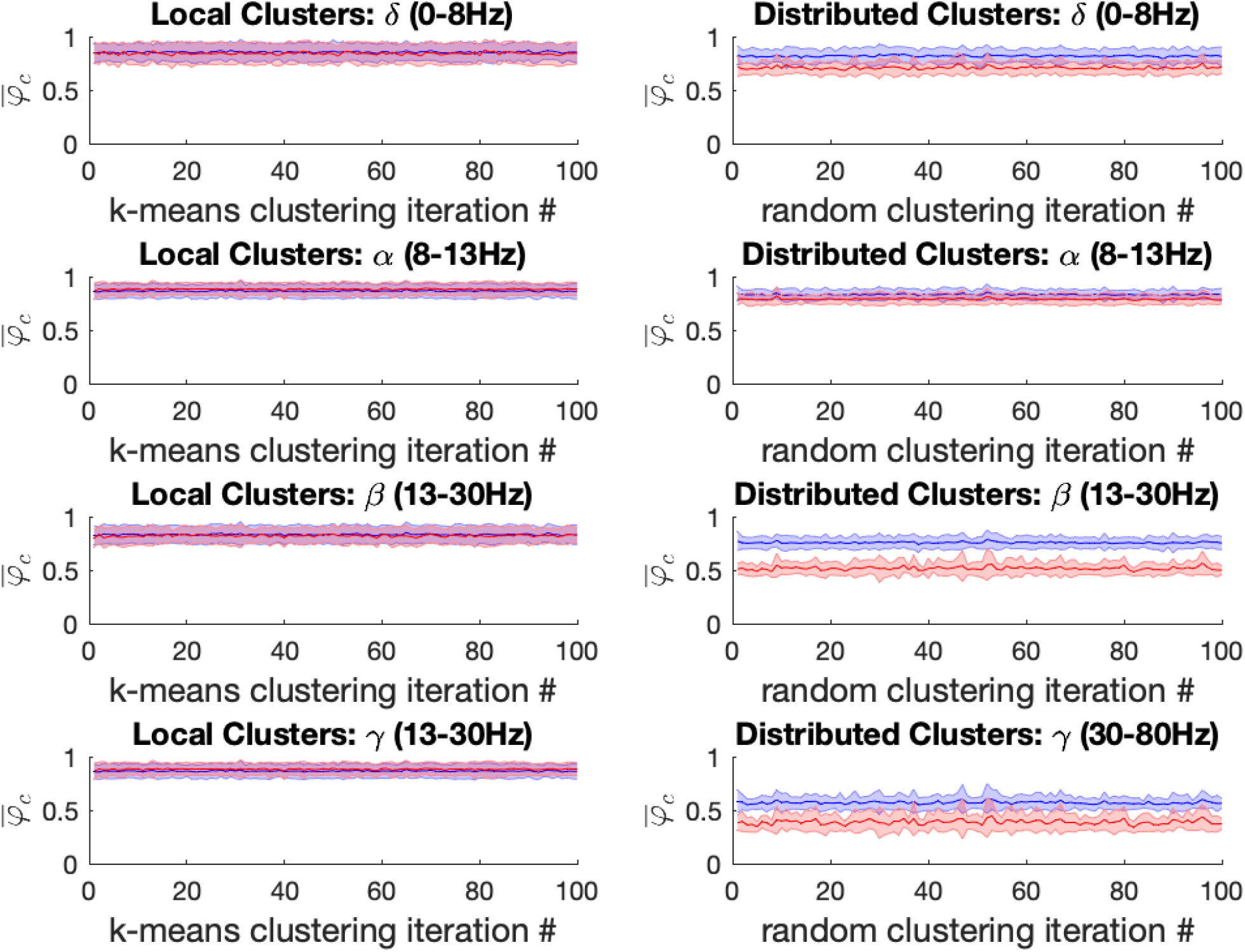
Left: For the propofol (red) vs baseline (blue) conditions, the mean local cluster synchrony 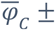 standard deviation is plotted for each of 100 k-means iterations (x-axis) in each of the four frequency bands under study for k = 13. Repeated within-condition t-tests failed to reach statistical significance at the Bonferroni-corrected threshold (α = 5*10^−4^) in any of the four frequency bands of interest. Right: Mean distributed cluster synchrony 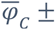 standard deviation is plotted for each of 100 iterations of the pseudo-random algorithm used for distributed cluster assignments (x-axis). Statistically significant reductions in distributed cluster synchrony at the Bonferroni-corrected significance threshold (α = 5*10^−4^) were observed following propofol anesthesia for all (100/100) distributed cluster assignments in the β and γ bands, respectively. By contrast only 3/100 significant differences in propofol’s effect on distributed cluster synchrony were observed in the α-band, while more mixed results were observed in the δ-band with 58/100 of distributed cluster assignments reaching significance.

**Supplementary 4.**
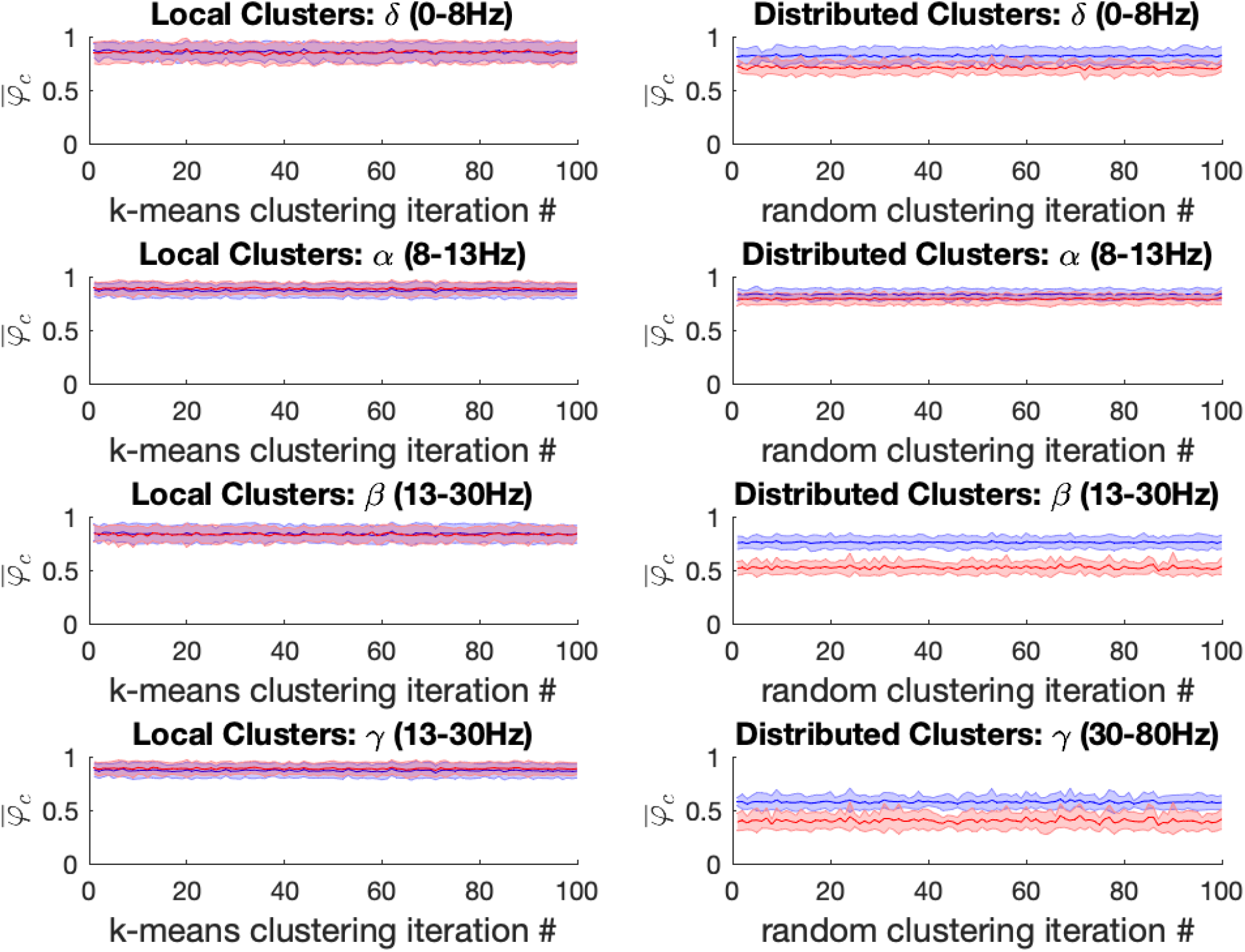
Left: For the propofol (red) vs baseline (blue) conditions, the mean local cluster synchrony 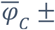 standard deviation is plotted for each of 100 k-means iterations (x-axis) in each of the four frequency bands under study for k = 15. Repeated within-condition t-tests failed to reach statistical significance at the Bonferroni-corrected threshold (α = 5*10^−4^) in any of the four frequency bands of interest. Right: Mean distributed cluster synchrony 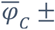 standard deviation is plotted for each of 100 iterations of the pseudo-random algorithm used for distributed cluster assignments (x-axis). Statistically significant reductions in distributed cluster synchrony at the Bonferroni-corrected significance threshold (α = 5*10^−4^) were observed following propofol anesthesia for all (100/100) distributed cluster assignments in the β and γ bands, respectively. By contrast only 7/100 significant differences in propofol’s effect on distributed cluster synchrony were observed in the α-band, while more mixed results were observed in the δ-band with 62/100 of distributed cluster assignments reaching significance.

**Supplementary 5.** Average p-values for the mean local cluster synchrony between the baseline and Ketamine conditions for the four-frequency bands across the the 100 k-means iterations for *k*=13,14 and 15. Significance is attained for the α band in 1 or two trial, but the average p-values do not reach significance at the Bonferroni-corrected significance threshold (α = 5*10^−4^).

|  | k=13 | k=14 | k=15 |
| --- | --- | --- | --- |
| $\delta$ | 0.634/(0/100) | 0.618/(0/100) | 0.641/(0/100) |
| $\alpha$ | 0.013/(1/100) | 0.010/(1/100) | 0.013/(2/100) |
| $\beta$ | 0.097/(0/100) | 0.019/(0/100) | 0.119/(0/100) |
| $\gamma$ | 0.358/(0/100) | 0.383/(0/100) | 0.397/(0/100) |

**Supplementary 6.** Average p-values for the mean distributed cluster synchrony between the baseline and Ketamine conditions for the four-frequency bands across the 100 pseudo-random distributed class assignment algorithm iterations for *k*=13,14 and 15. Significance (in bold) is attained for the α, β and γ band at the Bonferroni-corrected significance threshold (α = 5*10^−4^).

|  | k=13 | k=14 | k=15 |
| --- | --- | --- | --- |
| $\delta$ | 0.087/(0/100) | 0.09/(0/100) | 0.093/(1/100) |
| $\alpha$ | **$1*10^{-6}$/(100/100)** | **$3*10^{-7}$/(100/100)** | **$3*10^{-7}$/(100/100)** |
| $\beta$ | **$3*10^{-6}$/(100/100)** | **$8*10^{-7}$/(100/100)** | **$3*10^{-6}$/(100/100)** |
| $\gamma$ | **$1*10^{-5}$/(99/100)** | **$3*10^{-6}$/(100/100)** | **$2*10^{-6}$/(100/100)** |

**Supplementary 7.** Average p-values for the mean local cluster synchrony between the baseline and Propofol conditions for the four-frequency bands across the the 100 k-means iterations for *k*=13,14 and 15. Significance is never attained for any trial, and the average p-values do not reach significance at the Bonferroni-corrected significance threshold (α = 5*10^−4^).

|  | k=13 | k=14 | k=15 |
| --- | --- | --- | --- |
| $\delta$ | 0.695/(0/100) | 0.77/(0/100) | 0.769/(0/100) |
| $\alpha$ | 0.357/(0/100) | 0.28/(0/100) | 0.269/(0/100) |
| $\beta$ | 0.689/(0/100) | 0.74/(0/100) | 0.768/(0/100) |
| $\gamma$ | 0.816/(0/100) | 0.76/(0/100) | 0.744/(0/100) |

**Supplementary 8.** Average p-values for the mean distributed cluster synchrony between the baseline and Propofol conditions for the four-frequency bands across the 100 pseudo-random distributed class assignment algorithm iterations for *k*=13,14 and 15. Significance (in bold) is attained for the β and γ band at the Bonferroni-corrected significance threshold (α = 5*10^−4^).

|  | k=13 | k=14 | k=15 |
| --- | --- | --- | --- |
| $\delta$ | 0.006/(58/100) | 0.001/(60/100) | 0.003/(62/100) |
| $\alpha$ | 0.022/(3/100) | 0.017/(6/100) | 0.018/(7/100) |
| $\beta$ | $4 \cdot 10^{-7} / (100/100)$ | $1 \cdot 10^{-7} / (100/100)$ | $1 \cdot 10^{-6} / (100/100)$ |
| $\gamma$ | $2 \cdot 10^{-5} / (100/100)$ | $2 \cdot 10^{-5} / (100/100)$ | $1 \cdot 10^{-5} / (100/100)$ |

